# Admixture can readily lead to the formation of supergenes

**DOI:** 10.1101/2020.11.19.389577

**Authors:** Paul Jay, Thomas G. Aubier, Mathieu Joron

## Abstract

Supergenes are genetic architectures allowing the segregation of alternative combinations of alleles underlying complex phenotypes. The co-segregation of sets of alleles at linked loci is determined by polymorphic chromosomal rearrangements suppressing recombination locally. Supergenes are involved in many complex polymorphisms, including sexual, color or behavioral polymorphisms in numerous plants, fungi, mammals, fish, and insects. Despite a long history of empirical and theoretical research, the genetic origin of supergenes remains poorly understood. Here, using a population genetic two-island model, we explore how the evolution of overdominant chromosomal inversions may lead to the formation of supergenes. We show that the evolution of inversions in differentiated populations connected by gene flow leads to an increase in frequency of poorly adapted, immigrant haplotypes. When inversions are associated with recessive fitness cost hampering their fixation (such as a mutational load), this results in the formation of supergenes. These results provide a realistic scenario for the evolution of supergenes and inversion polymorphisms in general, and bring new light into the importance of admixture in the formation of new genetic architectures.

---

Discrete polymorphisms are widespread in nature. Besides the ubiquitous polymorphism of sexual and mating types (1, 2), classic examples of discrete polymorphisms involving multiple functional loci include polymorphic immune systems in vertebrates (3–5), and color polymorphisms in several birds (6–8), insects (9, 10), snails (11, 12) and lizards (13). Supergenes are genetic architectures which are often found to underlie discrete polymorphisms by preventing the formation of intermediate maladapted phenotypes. Supergenes are generally formed by polymorphic chromosomal rearrangements that suppress recombination among linked loci. Suppressed recombination is thought to facilitate the evolution of alternative combinations of co-adapted alleles (14). Such combinations are assumed to evolve as a result of selection favoring alternative evolutionary strategies (15).

The maintenance of polymorphism is a key feature of supergenes. Yet one would expect that the best evolutionary strategy, determined by a combination of alleles (i.e. a haplotype), should eventually fix in populations via selection. In nature, supergene alleles have been shown to be maintained by balancing selection, in particular by negative frequencydependent selection taking the form of disassortative mate choice (16) or heterozygote advantage (7, 8, 17, 18). Indeed, supergene alleles are often associated with chromosomal inversions that seem to harbour recessive deleterious mutations, as suggested by the lack of inversions homozygotes in natural populations (6, 8, 16, 18, 19). Inversions suppress recombination in heterozygotes, and are therefore prone to capture and accumulate deleterious mutations (18, 20, 21). This recessive mutational load may generate balancing selection and contribute to polymorphism at supergenes (18). Nevertheless, while the maintenance of supergene alleles is theoretically well understood, how these differentiated functional haplotypes arise remains a challenging question (22–25)

The difficulty of solving the origin of supergenes can be appreciated using the metaphor of the fitness landscape introduced by Wright (26). A population harbouring a supergene can be seen as a population with several non-recombining haplotypes underlying distinct phenotypes corresponding to alternative fitness peaks. The separation of fitness peaks in this model implies the existence of epistatic interactions; supergene haplotypes link together co-adapted alleles, and recombinant haplotypes thus display poor fitness (15, 23, 24, 27). To reach this situation, a polymorphic population may have evolved from a monomorphic populations. This implies a peak shift, across a fitness valley, forming differentiated haplotypes which do not reach fixation and thus segregate under balancing selection with an ancestral form. Peak shifts may occur via genetic drift (‘shifting balance theory’ (26)), via varying selective conditions (28), via major effect mutations (24), or when mutation rate and population size are high (29). Nevertheless, the processes underlying peak shifts do not explain the maintenance of ancestral and derived adaptive strategies in a polymorphism.

The puzzle of the origin and the maintenance of polymorphisms at supergenes rests on the antagonistic effects of recombination on the evolution of new combination of coadapted alleles. On the one hand, recombination can promote peak shifts by bringing into linkage beneficial mutations that arose on different chromatids. On the other hand, recombination can mix up alternative combinations of alleles and thus hamper the formation of new differentiated haplotypes (30). Therefore, recombination can promote the evolution of new adaptive strategies, but is also a homogenizing force that inhibits their maintenance in a polymorphism (24, 31). As a corollary, reduced recombination, caused by chromosomal rearrangements or by tight linkage between adaptive loci, can allow the maintenance of alternative haplotypes, but may also prevent peak shift and the formation of differentiated haplotypes.

Because of this dual effect of recombination, polymorphism is unlikely to be maintained during the process of peak shift (29). Nonetheless, the evolution of supergene haplotypes may occur in situations where rearrangements reducing re-combination occur after peak shifts. Indeed, a solution to this paradox may involve the evolution of alternative supergene haplotypes in either (i) different genomic regions or (ii) different taxa (14, 32, 33). In this later case, haplotypes belonging to the same population recombine normally, but not with haplotypes from the other population. Alternative strategies encoded by differentiated haplotypes can readily evolve in separated populations if they experience variation in selective pressure, and the same rationale applies for alternative haplotypes evolving in distinct genomic regions, for instance following a duplication. By the way of genomic translocation or migration between population (admixture), the different haplotypes may be grouped respectively (i) in the same genomic region or (ii) in the same population. If these haplotypes are not recombining, for instance because one of them has experienced a chromosomal inversion, and if some form of balancing selection maintains them in coexistence, they may form a supergene.

The scenario of genomic translocation has received low empirical support (but see (34)), but the admixture scenario has recently been shown to be involved in the formation of a wing-pattern mimicry supergene in the butterfly *Heliconius numata* (35), in the formation of a new sex-chromosome in the ninespine stickleback, *Pungitius pungitius* (36), and is suspected in several other cases (e.g. (8, 17)). The formation of a supergene by admixture involves migration between populations, evolution of suppressed recombination via chromosomal rearrangements, co-adaptation at several epistatic loci with dominance, and a force generating balancing selection. The interaction between these processes may generate complex evolutionary dynamics and, therefore, the conditions favoring the formation of supergenes are still obscure. Previous theory has focused on understanding the conditions favoring non recombining, rearranged haplotypes, demonstrating that they should spread in a broad range of genetic and demographic situations when they reduce the recombination load generated by migration (33, 37–44). Notably, Kirkpatrick and Barton (2006) (42) have shown that inversions can spread when they capture several beneficial mutations, but that inversions carrying recessive fitness costs cannot fix in populations and are compelled to segregate at intermediate frequency. We hypothesize, therefore, that under some conditions, a stable polymorphism could be formed by an inverted locally-adapted haplotype and a standard haplotypes (brought by migration), which could provide a general model for supergene evolution.

Here we construct a diploid two-locus population genetic model to investigate how a supergene can evolve when adjacent populations experiments different selection regimes. We consider the occurrence of a recombination modifier acting on heterokaryotypes such as a chromosomal inversion and associated with a fitness cost when homozygous, which generate overdominance and thus balancing selection. By tracking the frequencies not only of inverted segments but of all allelic combinations in a two population model, our study provides new insight into the causes and consequences of the spread of inversions. Using stochastic simulations, we show that under a wide diversity of fitness landscapes, gene flow between population is likely to lead to the formation of a supergene, suggesting that admixture may play an important role in the formation of new genetic architectures. In particular, we show that the formation of a supergene strongly depends on the interaction between frequency-dependent effects of recombination and of overdominance.

## Methods

### Purpose

We investigate the conditions under which chromosomal inversions spread in neighboring populations under disruptive selection, and how it associates with the formation of supergenes underlying the maintenance of discrete polymorphisms. We define a supergene as a genetic architecture maintaining under balancing selection multiple nonrecombining haplotypes encoding alternative discrete phenotypes. Polymorphisms at supergene is thus maintained over the long term by evolutionary forces balancing haplotype frequencies. We do not consider that polymorphisms balanced by migration-selection only are controlled by supergenes. Thereby, we aimed to study the formation of polymorphisms maintained by balancing selection in isolated populations, i.e. that can persist in the absence of a flow of alternative haplotypes from a neighbouring population.

Using a standard stochastic diploid population genetic model, we implement a chromosomal rearrangement as a modifier allele that controls the recombination rate between two ecological loci. The expression of this modifier allele at homozygous state may also associate with a fitness cost that reflects the deleterious effect of mutation accumulation in chromosomal inversions (21).

Simulations are divided into two phases. First, populations evolve under migration-selection-mutation balance (migratory phase). Second, populations keep evolving but this time under a selection-mutation balance (non-migratory phase). With such simulation experiments, we aim at describing in detail the forces maintaining polymorphism in populations and the conditions under which a discrete polymorphism is formed and stably maintained even without migration.

### Spatial structure and genotypes

We model a stochastic system of two diploid populations exchanging migrants (two-island model). This population genetic model postulates three autosomal diploid loci with alternative alleles represented by small vs capital letters. Two ecological loci, A and B, are subject to disruptive selection and can recombine. The third locus, M, is physically linked to locus A (no recombination) and can impede recombination among loci A and B (Fig. 1). More specifically, in heterozygotes with genotype *Mm* at the M locus, there is no recombination between loci A and B. The invasion of allele *M* at the modifier locus reflects the invasion of a chromosomal inversion, and we therefore denote the presence of the recombination modifier in a genotype by inverting the order of loci. For instance, the haplotype *m* − *A*− *B* is referred to as *AB*, and the haplotype *M* − *A* − *B* is referred to as *BA* (Fig. 1). We refer to haplotypes with the *M* allele as ‘inverted’ haplotypes and haplotype with *m* allele as ‘ancestral’ haplotypes.

**Fig. 1.**
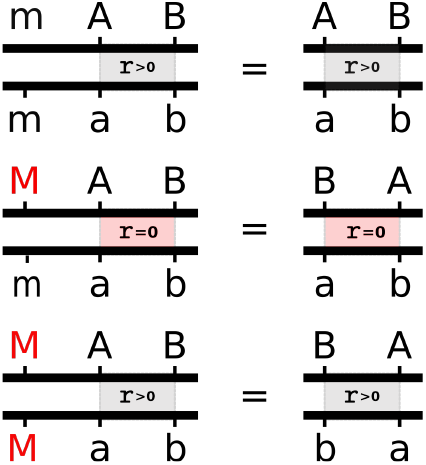
Examples of genotypes in the population genetic model (left), and equivalence in term of chromosomal rearrangements (right). Inverted and ancestral haplotypes cannot recombine (i.e., recombination rate *r* = 0 if the genotype at the M locus is *Mm*; middle row).

There are thus 2^3^ = 8 possible combinations of alleles that can be found on any chromosome (i.e., haplotypes): *mAB, mAb, maB, mab, MAB, MAb, M aB* and *M ab*. In each population *p* ∈ {1, 2}, the frequency of each haplotype *i* ∈ {1, …, 8} of alleles is referred to as *x*_*p,i*_.

Each individual carries two copies of the chromosome. As a result, our system is characterized by 8 *×* 8 = 64 genotypes. In each population *p* ∈ {1, 2}, the frequency of each genotype *j* ∈ {1, …, 64} deriving from those unions is referred to as *y*_*p,j*_. Assuming discrete generations, we follow the evolution of genotype frequencies **y**(*t*) within a finite population over time. **y** = {*y*_*p,j*_} is a vector consisting of 128 elements referring to the frequencies of the 64 genotypes present in newborn offspring in the two populations. Haplotype frequencies *x*_*p,i*_ are easily be calculated from **y**.

### Stochastic simulations

#### Migration

The life cycle begins with symmetric migration between populations at rate *m*, which describes the proportion of each population that consists of migrants right after the migration event. Thus, the frequency of genotype *j* in population *p* is:

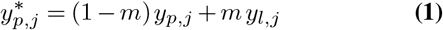

where *l* = 2 when *p* = 1, and *l* = 1 when *p* = 2

#### Selection

Migration is followed by natural selection in each population. The relative fitness associated with genotype *j* in population *p* is *w*_*p,j*_ ∈ [0, 1]. After natural selection, genotypic frequencies in population *p* are therefore:

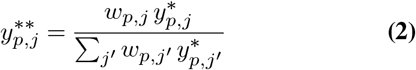

The nature of the fitness landscape in each population (parameter values *w*_*p,j*_) are detailed below in subsection ‘Simulation experiments’.

### Recombination, mating and zygote formation

At this point, gametes are produced. Depending on the genotype at the modifier locus M, recombination between loci A and B may occur during gamete production. The recombination rate between loci A and B is equal to 0 for a genotype *Mm* at the M locus, and is equal to *r* for genotypes *MM* and *mm* (Fig. 1). In each population, mating is random. In the new generation, expected genotype frequencies 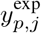 of zygotes are therefore calculated by summing the appropriate mating frequencies, assuming Mendelian segregation and accounting for recombination between loci A and B.

### Stochastic sampling and mutations

The new vector **y**(*t* + 1) is obtained by randomly sampling *N* offspring individuals from the expected genotype frequencies 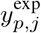 of zygotes. Additionally, in each offspring individual, we assume that mutation can occur with a probability *µ* per allele at loci A and B. We also assume that at locus M, allele *m* can mutate to allele *M* with a probability *µ*_inv_. This reflects the rate of inversions occurrence. Since such rearrangements are unlikely to affect the same portion of the genome twice, we assume unidirectional mutation at locus M.

### Simulation experiments

#### Fitness landscape

The phenotype expressed by each genotype *j* determines its fitness *w*_*p,j*_ in each environment (i.e., in population *p*). Phenotypes expressed at loci A and B are [*AB*], [*Ab*], [*aB*] and [*ab*]. Both environments have two peaks of fitness corresponding to phenotype [*AB*] and [*ab*]. The main peak being [*AB*] in population 1 and [*ab*] in population 2, and the second peak being [*ab*] in population 1 and [*AB*] in population 2 (Fig. 2). We consider a symmetric fitness landscape, such as:

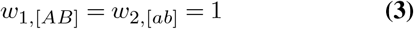

and

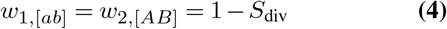

where *w*_*p*,[*AB*]_ (resp. *w*_*p*,[*ab*]_) corresponds to the fitness associated with phenotype [*AB*] (resp. [*ab*]) in population *p*, and where *S*_div_ reflects the fitness difference between the two local peaks. We refer to the best haplotype in a population as the ‘local’ haplotype (*AB* in population 1 and *ab* in population 2), and the haplotype corresponding to the second fitness peak as the ‘immigrant’ haplotype, i.e the haplotype that is favored in the neighboring population, even if this haplotype can eventually be maintained in the local population without gene flow. For instance, *ab* is considered as the ‘immigrant’ haplotype in population 1. We referred to *Ab* and *aB* haplotypes as ‘recombinant’ haplotypes. These haplotypes are associated with phenotypes [*Ab*] and [*aB*], considered ‘Valley’ phenotypes, such that:

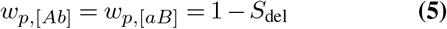

where *S*_del_ reflects the reduction in fitness associated with these intermediate valley phenotypes. We implement fitness landscapes such that 0 *<* 1 − *S*_del_ *< -* 1 *S*_div_ *<* 1 (Fig. 2).

**Fig. 2.**
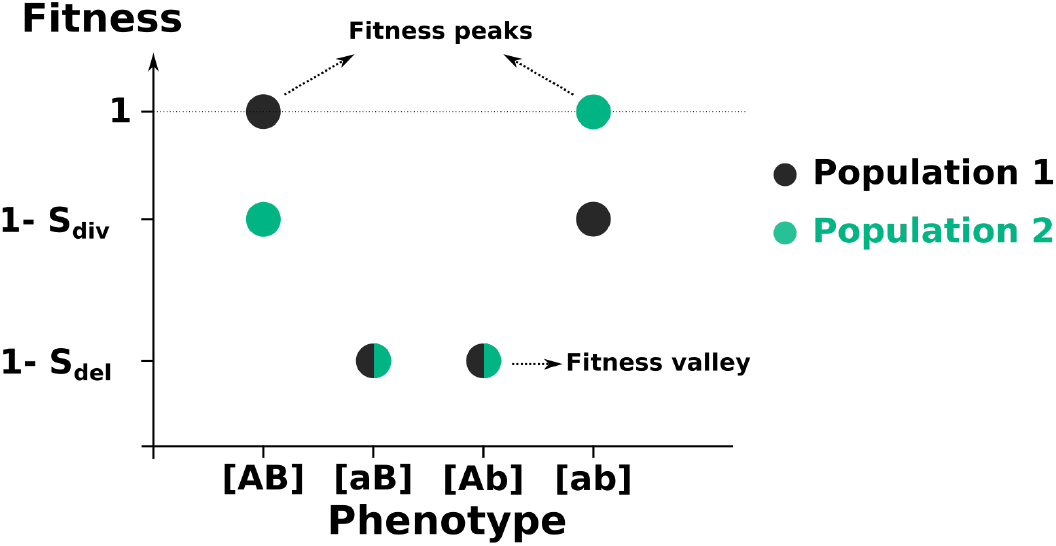
Representation of the fitness landscape implemented.

For heterozygous genotypes, we assume that dominant phenotypes (carrying alleles *A* and *B*, i.e., represented by capital letters) are expressed proportionally to a dominance factor *h* ∈ [0.5, 1]. For instance, the fitness associated with genotype *Ab/AB* in environment *p* is equal to:

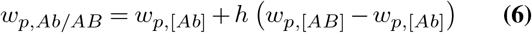

We refer to *AB* haplotypes as ‘dominant’ haplotypes (which is the best haplotype in population 1), and *ab* haplotypes as ‘recessive’ haplotypes (which is the best haplotype in population 2).

#### Fitness cost associated to *MM* homozygotes

The allele *M* at the M locus may correspond to rearrangements like chromosomal inversions that may carry recessive deleterious mutations (20). The expression of those deleterious mutations at homozygous state are thus associated with fitness cost. Therefore, we assume that the fitness of homozygotes *MM* is multiplied by a factor 1 −*γ*_inv_. For instance

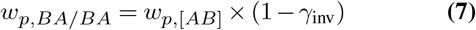

Parameter *γ*_inv_ ∈ [0, 1] reflects the fitness cost in homozygotes at the M locus. For *γ*_inv_ = 0, *MM* homozygotes do not incur a fitness cost. For *γ*_inv_ = 1, *MM* homozygotes incur a strong fitness cost; they all die during the selection process. Since it affects only homozygotes, this fitness cost generates overdominance: heterozygotes perform better than homozygotes.

#### Initialization

Simulations start with only *AB* and *ab* homozygotes carrying alleles *m* at the M locus in the two populations.

#### Simulations

Each simulation is divided into two phases, each lasting 5,000 generations. During the first phase, migration occurs at a rate 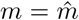 Then, during the second phase, there is no migration between the two populations – i.e., *m* = 0. Those phases are referred to as ‘migratory phase’ and ‘non-migratory phase’, respectively. The objective of the non-migratory phase is to estimate to what extent the polymorphism formed in migratory phase depends on the migration. Thereby, we aimed to study polymorphisms not only maintained by migration-selection balance but also by balancing selection in isolated populations.

Table 1 sums up parameters notations and provides the list of values implemented for each parameter in our simulations. We performed 10 simulations for each of the 34,020,000 combinations of parameter values tested.

**Table 1.** Summary of parameters implemented in simulations.

| Parameter | Definition | Values |
| --- | --- | --- |
| $\hat{m}$ | Migration between populations during the migratory phase | { 0.05, 0.10, 0.15 } |
| $r$ | Recombination rate between loci A and B | { 0.0, 0.1, 0.2, 0.3, 0.4, 0.5 } |
| $\gamma_{inv}$ | Inversion recessive load | { 0.00, 0.05,... , 0.95, 1.00 } |
| $S_{div}$ | Difference in fitness between local and immigrant haplotypes | { 0.2, 0.3,... , 0.8, 0.9 } |
| $S_{del}$ | Difference in fitness between local and recombinant haplotypes | { 0.0, 0.1, 0.2, 0.3, 0.4, 0.5 } |
| $\mu$ | Mutation rate | { $10^{-8}$, $10^{-7}$, $10^{-6}$, $10^{-5}$, $10^{-4}$ } |
| $\mu_{inv}$ | Inversion rate | { $10^{-8}$, $10^{-7}$, $10^{-6}$, $10^{-5}$, $10^{-4}$ } |
| $N$ | Population size | { $10^3$, $10^4$, $10^5$ } |
| $h$ | Dominance level | { 0.50, 0.75, 1.00 } |

We found that the mutation rate and the inversion rate have little influence on the simulations, as long as they occur several times during the simulation times. We therefore show the results with a mutation rate equal to 10^−6^and a recombination modifier mutation rate equal to 10^−6^, which means that a recombination modifier is expected to arise around 50 times in a population of size *N* =10,000 during 5,000 generations.

## Results

Conducting simulations under a wide range of parameter values, we observed seven distinct scenarios regarding the fate of newly evolved chromosomal inversions (Figs. 3 and S1-S2; the 7th scenarios is shown only in Fig. S2). Inversion may not spread at all (Fig. 3B). When the inversion spreads, either on the dominant or the recessive haplotypes, it can fix (Fig. 3C-D) or be maintained at intermediate frequency during the migratory phase (Fig. 3E-G). This polymorphism can either be lost during the non-migratory phase (Fig. 3E-F) or be maintained (Fig. 3G). In this later case, the polymorphism involves inverted local haplotypes and ancestral immigrant haplotypes, and can be thus considered to be controlled by a supergenes. Simulations conducted using different life cycles resulted in the same scenarios in similar proportion (Fig. S3-S4).

**Fig. 3.**
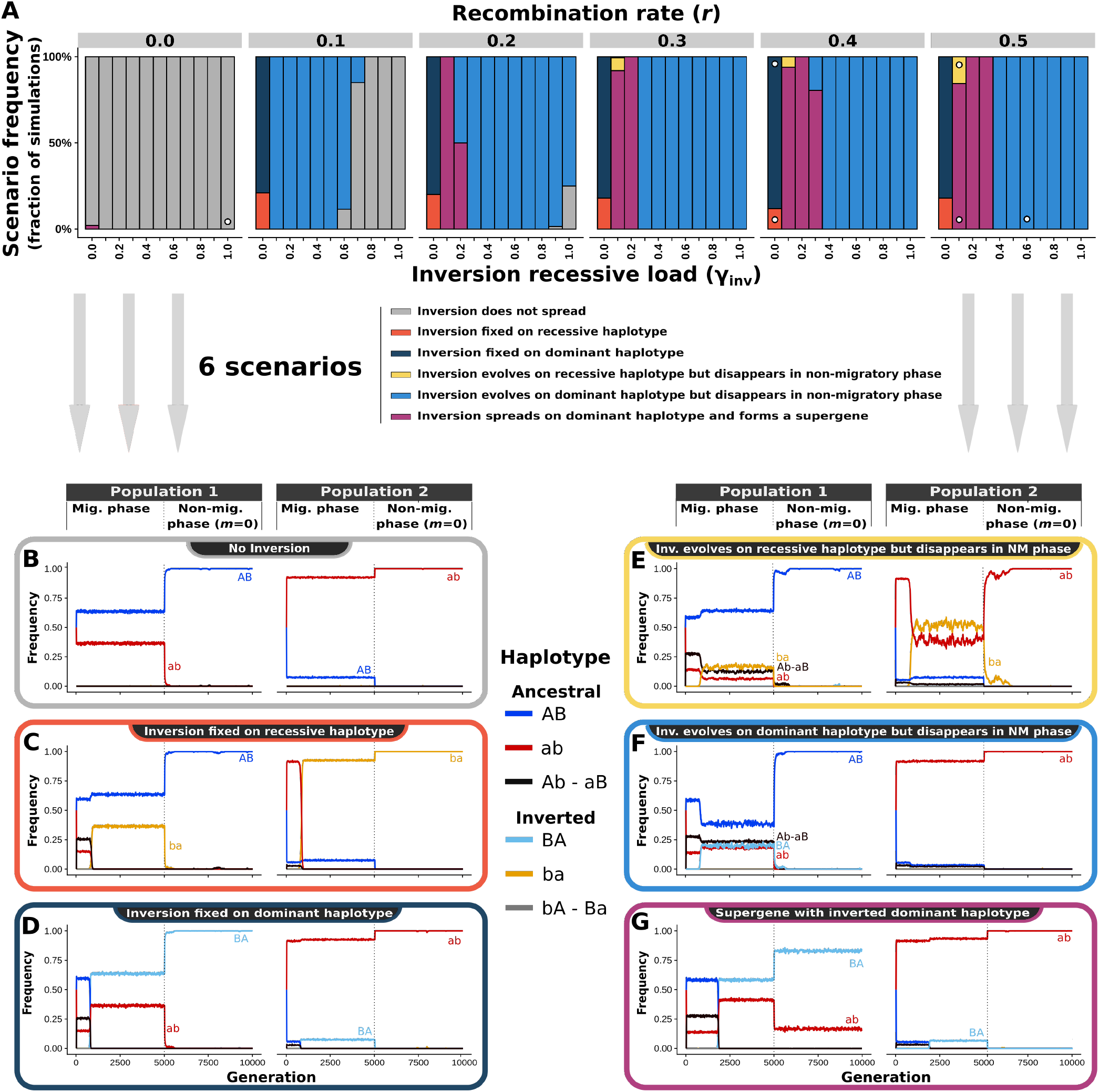
The evolution of chromosomal inversions, and consequences for the formation of supergenes. Simulations lead to six possible scenarios, depending on migration rate, recombination rate, and the strength of the inversion recessive load. **A** Effect of inversion recessive load and recombination rate on the fate of chromosomal inversions. Parameters used: 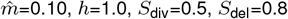. White dots indicate parameters used for each scenario plots (**B**-**G**). **B** Inversions do not spread. **C** Inversions fix, associated with recessive haplotypes (in population 2). **D** Inversions fix, associated with dominant haplotypes (in population 1). **E** Inversions evolve in association with recessive haplotypes, segregate at intermediate frequency with ancestral dominant and recessive haplotypes during the migratory phase, but disappear during the nonmigratory phase. **F** Inversions evolve in association with dominant haplotypes, segregate at intermediate frequency with both ancestral haplotypes during the migratory phase, but disappear during the non-migratory phase. **G** Inversions evolve in association with dominant haplotypes and segregate at intermediate frequency with only the ancestral recessive haplotypes in both migratory and non-migratory phases.

### Conditions for the spread of an inversion

In simulations where loci *A* and *B* never recombine, inversions do not associate with any benefit, and therefore rarely spread (Fig. 3A; e.g. Fig. 3B). When recombination is low and the recessive load carried by inversions is high, the low benefit of reducing recombination is offset by the cost brought by the recessive load, and inversions do not spread (Fig. 3A). Finally, a low fitness difference between local and immigrant haplotypes leads to swamping by dominant alleles (alleles A and B invade in both populations, Fig. S5). Therefore, in the absence of allelic diversity, suppressing recombination bears no advantage and newly evolved inversions do not increase in frequency. Overall, inversions reducing the recombination between coadapted alleles can easily spread in the population when they associate with a recessive load relative to the recombination rate among captured loci (all scenarios that are not colored in gray in Fig. 3A)(33, 42).

### Condition for the fixation of an inversion

Natural selection can indirectly favor the spread of inversions. Inversions suppress recombination and thereby reduce the fraction of offspring with a valley phenotype (Fig. 4B). This effect is observed in the two populations, whatever the haplotype captured by the inversions (Fig. S6). When inversions capture two co-adapted recombining loci, inverted haplotypes rise in frequency and tend to replace ancestral haplotypes (Figs. 3 and 4). Nevertheless, ancestral local and ancestral immigrant haplotypes are not affected in the same way (Fig. 4A-C). Immigrant haplotypes segregate at lower frequency than local haplotypes, and consequently they recombine more. Perhaps surprisingly, the rise of inversions on local haplotypes is therefore beneficial to immigrant haplotypes because it decreases the frequency of haplotypes they recombine with. The frequency of immigrant haplotypes therefore increases when inversions spread (Figs. 3C-G and 4C), and this occurs whatever the immigrant fitness (as long as there is no gene swamping, Fig. S8). Conversely, ancestral local haplotypes do not benefit from this effect and are not constantly brought by migration, and thus tend to disappear when inversions reach high frequency (Figs. 3C-G and 4A).

**Fig. 4.**
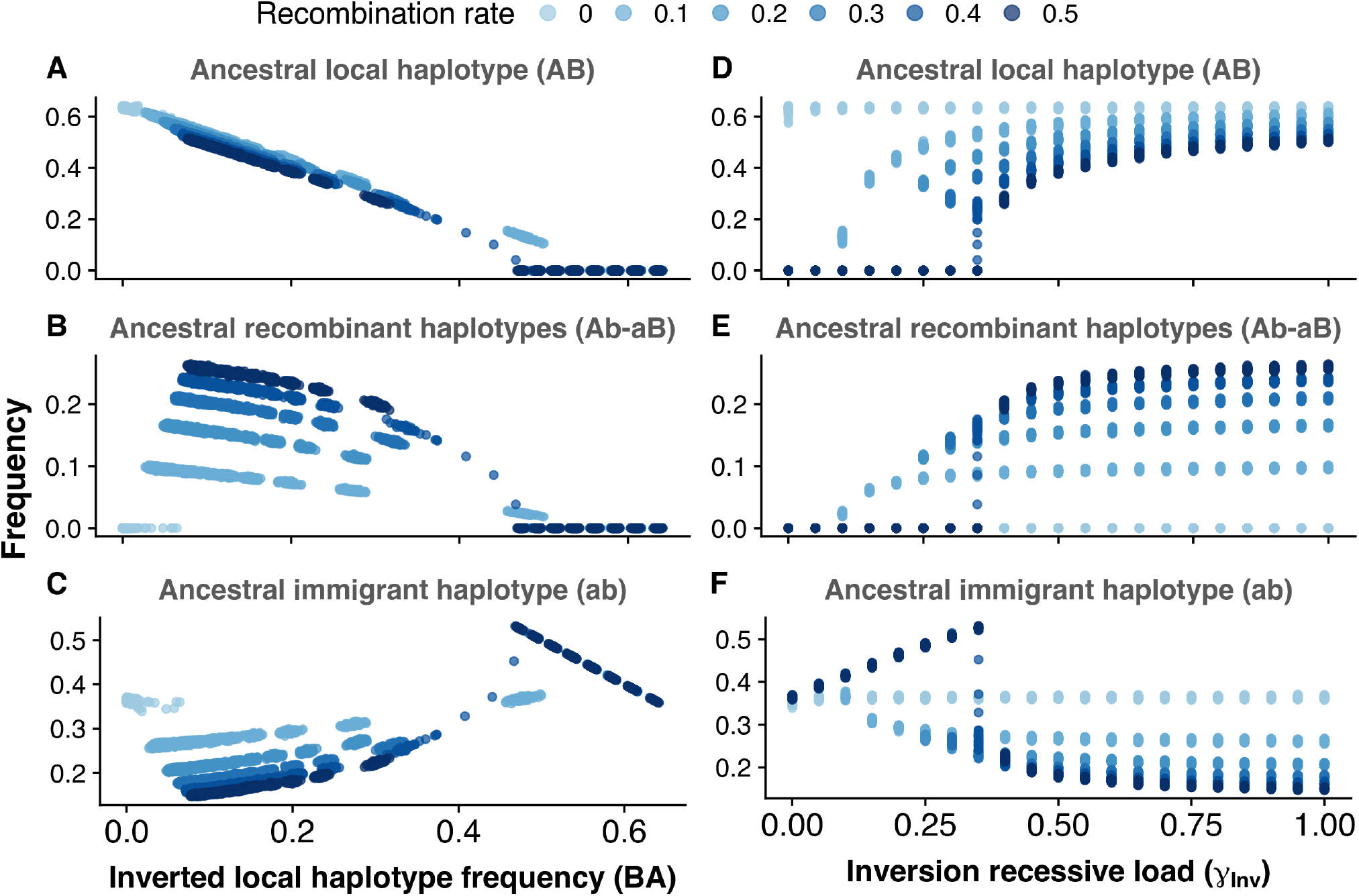
Consequences of the evolution of inversions on the frequencies of ancestral haplotypes. Plots display the results for the spread of inversions capturing the dominant haplotype in population 1, since inversions on the recessive haplotype are less likely to spread and to reach high frequencies (in population 2). **A** Effect of the spread of inverted haplotypes on the frequency of ancestral local haplotypes. **B** Effect of the spread of inverted haplotypes on the frequency of recombinant haplotypes. **C** Effect of the spread of inverted haplotypes on the frequency of ancestral immigrant haplotype. **D** Effect of the recessive fitness cost associated with an inversion on the frequency of ancestral local haplotypes **E** Effect of the recessive fitness cost associated with an inversion on the frequency of recombinant haplotypes. **F** Effect of the recessive fitness cost associated with an inversion on the frequency of ancestral immigrant haplotypes. Parameter values used for these analyses : 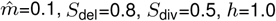.

Overall, when inversions are associated with a low recessive load (*γ*_inv_), they tend to replace their ancestral counterparts and segregate with ancestral immigrant haplotypes at migration-selection balance during the migratory phase (Fig. 3C-D). When migration stops, immigrant haplotypes are rapidly eliminated by selection and inversions reach fixation (Fig. 3A-C-D). The situation is rather different if the inversion associates with an intermediate recessive load, as detailed below.

### The fate of overdominant inversions

When inversions associate with an intermediate recessive load (*γ*_inv_), inversions behave as overdominant and cannot fix in the populations. Recessive deleterious effects expressed by homozygous inversion genotypes indeed generate heterozygote advantage, which translates into negative frequency-dependent selection acting on inversions. A recessive load generates nearly no cost when inversions are rare but become powerful when they reach higher frequencies. Therefore, increasing the inversion load *γ*_inv_ does not affect the probability of spread of inversions (Fig. S9) but determines their frequency at equilibrium (Fig. S11). As a consequence, *γ*_inv_ strongly determines their long-term fate (Figs. 3A and S11).

The overdominant behavior of inversions makes them prone to form a polymorphism with ancestral haplotypes and under certain conditions, this polymorphism persists in non-migratory phase and can be considered as forming a super-gene. Two majors behaviors can indeed be observed when overdominant inversions evolve:

### When overdominant inversions do not replace their ancestral equivalent

As described above, the rise of inverted haplotypes leads to a decrease in the frequency of their ancestral counterpart and an increase in the frequency of the immigrant haplotypes. When inversions remain at low frequency, they do not replace their ancestral counterparts and this leads to a coexistence of inverted local, ancestral local and ancestral immigrant haplotypes during the migratory phase (Fig. 3E-F). During the non-migratory phase, immigrant haplotypes are rapidly lost, so inversions no longer bring a fitness advantage and are impaired by their recessive fitness loads. Consequently, when inverted (e.g. *BA*) and ancestral (e.g. *AB*) haplotypes segregate at end of the migratory phase, the ancestral form always fixes in the population as soon as migration is stopped and inversions are lost (Figs. 3E-F and S10). Notably, we get this outcome in all deterministic simulations without genetic drift (i.e., without stochastic sampling; not shown), as long as inverted haplotypes can spread initially.

### When overdominant inversions replace their ancestral equivalent

The rise of inversions at high frequency may lead to the crash of their ancestral equivalent but not of the ancestral immigrant haplotypes. When inversions are overdominant, however, they do not fix in the population during the non-migratory phase, because of negative frequency-dependent selection (Fig. S11). Therefore, when overdominant inversions have replaced their ancestral counterpart and segregate only with ancestral immigrant haplotypes at the end of the migratory phase, this polymorphism is stably maintained during the non-migratory phase (Figs. 3G and S10). Since this polymorphism is maintained by balancing selection and composed by non-recombining haplotypes coding for alternative phenotypes, the genetic architecture underlying this polymorphism can be considered a supergene.

Overall, the fate of an overdominant inversion during the non-migratory phase is therefore determined by whether its ancestral equivalent is lost or not during the migratory phase

### The determinants of overdominant inversion equilibrium frequency

Our simulations account for a wide diversity of demographic, genetic and selective landscapes. This allows us to highlight the factors controlling the frequency of overdominant inversion at a migration-selection equilibrium, which determines the persistence of polymorphisms in non-migratory phase.

First, the frequency reached by an inversion during the migratory phase strongly depends on the dominance of the haplotype captured by the inversion. Since heterozygotes benefit from a better fitness when dominance is strong (phenotype closer to the dominant fitness peak), inversions reach higher frequencies when they capture dominant haplotypes (Fig. S12). Recessive inversions are less likely to reach fixation than dominant inversions (for *γ*_inv_ = 0, Fig. 3A), and overdominant inversions never reach high frequency or form a protected polymorphism when capturing recessive haplotypes (Figs. 3A, S2 and S12). The spread of an inversion capturing one haplotype also inhibits the spread of an inversion capturing another haplotype (Fig. S13). Therefore, the dynamics of a simulation depend on which inversion spreads first, and this explains why, when the fitness load is low (e.g. *γ*_inv_ = 0.1), the formation of supergenes appears slightly less likely (Fig. 3A): by chance, an inversion capturing the recessive haplotype arises and spreads first, hampering the spread of other inversions capturing the dominant haplotype (Fig. 3E).

The major determinants of inversions frequency stand as the strength of their associated recessive load (*γ*_inv_), the migration rate 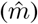 and the recombination rate (*r*) (Figs. 3A, S1 and S11). Since inversions are indirectly selected because they decrease the frequency of recombinant haplotypes resulting from recombination between local and immigrant haplotypes, inversions spread at higher frequency under high migration and recombination rates (Fig. S7). The recessive load generates a fitness cost on inversions that strengthens as they increase in frequency (because of the increasing frequency of homozygotes), and imposes an equilibrium frequency to inversions (Fig. S11). This leads to a dramatic change in frequency of ancestral and inverted haplotypes when this load increases (Figs. 4D-F and S11). The interaction between recombination rate, migration rate and recessive load thus determines if an overdominant inversion forms a protected polymorphism or disappears during the non-migratory phase.

When the recessive load is low relative to the recombination rate, inversions can easily spread (Figs. 3A, S1 and S2). When the recessive load is high relative to the recombination rate, the inverted haplotype never reaches a sufficient frequency to replace ancestral haplotypes, and thus the inversion is lost during the non-migratory phase (Fig. 3F). Nonetheless, for intermediate values, drift can lead to the loss of the ancestral haplotype, and to the formation of a supergene (as depicted in Fig. 3A for *r*=0.2 and *γ*_inv_=0.2).

Interestingly, the fitness associated with the valley phenotype as well as the immigrant phenotype appear as poor determinants of the frequency of inversions and the formation of supergenes (Figs. S14 and S15). This suggests that supergenes can be formed by haplotypes with large variation in fitness components and under a wide diversity of fitness landscapes.

## Discussion

Using a two-population model, we explored how a recessive fitness load associated with a chromosomal inversion influences the fate of this inversion and the formation of a protected discrete polymorphism at multiple linked loci. We show that recombination and the recessive fitness load have opposing effects on the frequency of inversion, which deterministically leads to the formation of a supergene when these forces are balanced. Our results, showing the effect of migration and recombination rates on the invasion of a chromosomal inversion, are consistent with Kirkpatrick and Barton’s (2006) analytical derivations (42), but by tracking the frequencies of all haplotypes in two diploid populations under different levels of (over)dominance, epistasis and recombination, we were also able to determine how the invasion of inversions affects the frequencies of ancestral (noninverted) haplotypes. Perhaps surprisingly, the spread of inversions does not only decrease the frequencies of local ancestral and recombinant haplotypes, but also increases the frequency of immigrant haplotypes irrespective of their local fitness and dominance levels. When the recombination rate is high, a moderate recessive fitness cost associated with inversions leads to the loss of local ancestral haplotypes as well as the maintenance of locally maladapted, immigrant haplotypes. This leads to the formation of a supergene characterized by the maintenance of a polymorphism with a locally adapted inverted haplotype and a locally maladapted standard haplotype, even though these haplotypes display large differences in their degree of local adaptation.

Here, we restricted our definition of supergenes to those cases of discrete polymorphisms that can be maintained in absence of gene flow. This strict definition allowed us to untangle the forces contributing to the emergence of inversion polymorphisms within populations. In natural populations, however, it is difficult to precisely assess the effect of migration among other factors. Yet we believe it is useful to recognise the differences between polymorphisms reflecting the influx of migrants from those owing to frequency-dependent processes within populations. As shown on Fig. 3, a continuum of situations can be observed, characterised by variations in the importance of migration in the maintenance of polymorphism. We consider the term supergene is most useful to describe the genetic architectures which facilitate the maintenance of polymorphism and the segregation of discrete forms (15), as in most of the classic cases on distyly, mimicry, shell morphologies, or behavioral syndromes (6, 8–11, 34, 45). Those tend to be cases with known or suspected epistasy among linked loci as well as some level of overdominance, and represent one end of the spectrum predicted by our model (case represented in Fig. 3F). Other well-described cases of inversion polymorphisms such as those distributed along clines or ecotones (46–48), might represent cases at the other end of the spectrum, where migration plays a large role in the maintenance of a high level of polymorphism (cases represented in Fig. 3C and 3D). Whether these inversion controlling the ecotypes distributed along environmental gradients also tend to harbour recessive deleterious variants that may contribute to their maintenance in the face of varying migration levels still remains to be elucidated. The major point made here is, however, that the evolution of supergenes maintained by local frequency-dependent selection on overdominant inversions may be facilitated by spatial segregation of selection.

Overdominance of inversions may have two major causes. First, inversion breakpoints may directly generate deleterious effects, for instance when they disrupt a gene. Second, recessive deleterious variants may be present in inversions, either captured during their formation, or subsequently accumulated due to recombination suppression (21). Our population genetic model treats the recombination load and the inversion recessive load as independent parameters. Yet, the size of an inversion determines both the potential number of captured deleterious mutations and the linkage of captured adaptive loci. Therefore, the fate of a chromosomal inversion – whether it invades a population, remains polymorphic or forms a supergene – may strongly depend on its size. This may explain why the frequency of inversion appears to correlate with their size in humans (49). Although we modelled the joint effect of recombination and mutation load on the spread of inversions, we did not model explicitly the size of the inversion, which links those two parameters and may lead to a trade-off: a large inversion is more likely to capture a favorable combination of recombining alleles but is also more likely to carry a high number of deleterious mutations. Intermediate-size inversions might be more prone to spread (and to form a supergene under some conditions) as a result of this putative trade-off. Further theoretical work is therefore required to understand how the fate of inversions can be affected by their size.

Overdominant inversions seem to be widespread in nature, and notably stand as a general explanation for supergene inheritance. For instance, in the social chromosome of the fire ant *Solenopsis invicta* (17) or in the supergene controlling plumage color and mating behaviour in the white-throated sparrow *Zonotrichia albicollis* (8), inversion homozygotes are not found in nature. Likewise, in the butterfly *Heliconius numata*, inversion homozygotes at the supergene controlling mimicry polymorphism display a very low larval survival (18) and are rare in nature (16). In these cases, the recessive load of inversions seems very high, whereas we predict that only low to moderate fitness cost can lead to the formation of supergenes. Because polymorphic inversions are expected to accumulate additional deleterious variants after their formation, these fitness cost may have strenghten after supergene formation.

Our model shows that negative frequency-dependent selection can favor the spread and the maintenance of supergene alleles with strong differences in fitness components. More specifically, an allele that is not locally favored can be maintained alongside a locally adapted allele if the latter is under negative frequency-dependent selection. This may explain why many supergenes are characterized by the maintenance of several phenotypes that seem to be associated with strong fitness differences. For instance, multiple-queen fire ant colonies seem to outcompete mono-queen colonies on many aspects but both strategies are maintained in nature (50). Likewise, mimetic and non-mimetic butterfly morphs coexists in several *Papilio* species while only mimetic morphs seem to benefit from reduced predation (51). Supergenes are often assumed to be maintained because environmental factors generate alternative adaptive optima that are associated with similar fitness consequences (14, 15). In contrast, our results suggest that supergenes could evolve because negative frequency-dependent selection caused by intrinsic features of chromosomal rearrangements leads to the maintenance of alternative haplotypes even if they are highly differentially adapted to the local environment.

Our model does not include major effects mutations and may apply in a wide diversity of demographic conditions. As such, it may offer a parsimonious explanation to the puzzle posed by the origin of differentiated supergene haplotypes. Considering separated taxa evolving in different conditions, differentiated haplotypes could easily evolve and be affected by chromosomal inversions independently. We show that gene flow between these taxa could result in the formation of supergenes. This scenario does not imply that these taxa are categorized as distinct species and the mechanism proposed here can readily apply to differentiated populations from the same species. During the past decade, progress in sequencing technologies have revealed that gene flow between differentially-adapted taxa is common in nature including between species (52, 53). Such gene flow may generate a recombination load, which fosters the spread of inversions or other chromosomal rearrangements which reduce recombination (42). This could explain why structural polymorphisms are widespread across the tree of life (e.g. (49, 54–56)). Our model could apply to several instance of supergene evolution, and indeed, several known supergenes display genomic signals suggesting an admixture scenario (8, 35, 36, 57).

Considering the pervasive nature of inversions in genomes, their capacity to capture deleterious variants, and the widespread gene flow among differentiated populations, our model shows that protected inversion polymorphisms can readily evolve and suggests that supergenes could be more frequent that previously thought. Supergenes can control the switch between distinct phenotypes at multidimensional traits that are not easily perceptible, such as variations in behavior (17, 58). Variation in sperm size and mobility in songbirds provides for instance a dramatic example of a recently discovered trait variation that is not easily observable but is controlled by a supergene (58). Known supergenes are nearly always associated with obvious color polymorphism (e.g. (7–9, 11, 12, 17); This is likely a detection bias and current advances in sequencing technology could thus reveal the presence of such architecture coding a wide diversity of traits in numerous species.

In summary, the recombination load generated by migration between differentiated populations is expected to foster the spread of recombination modifiers such as chromosomal inversions. Nevertheless, these recombination modifiers are prone to be associated with recessive fitness loads, which may translate into negative frequency-dependent selection hampering their fixation. This fosters the maintenance of polymorphism and could lead to the formation of supergenes. Taken together, theses results provide a realistic scenario for the evolution of supergenes and inversion polymorphisms in general and bring new light into the importance of admixture in the formation of new genetic architectures.

## Acknowledgements

This research was supported by grants from the Swiss National Science Foundation (to T.G.A.) and from the European Research Council (ERC-StG-243179) (to M.J.).

## Supplementary material

**Fig. S1.**
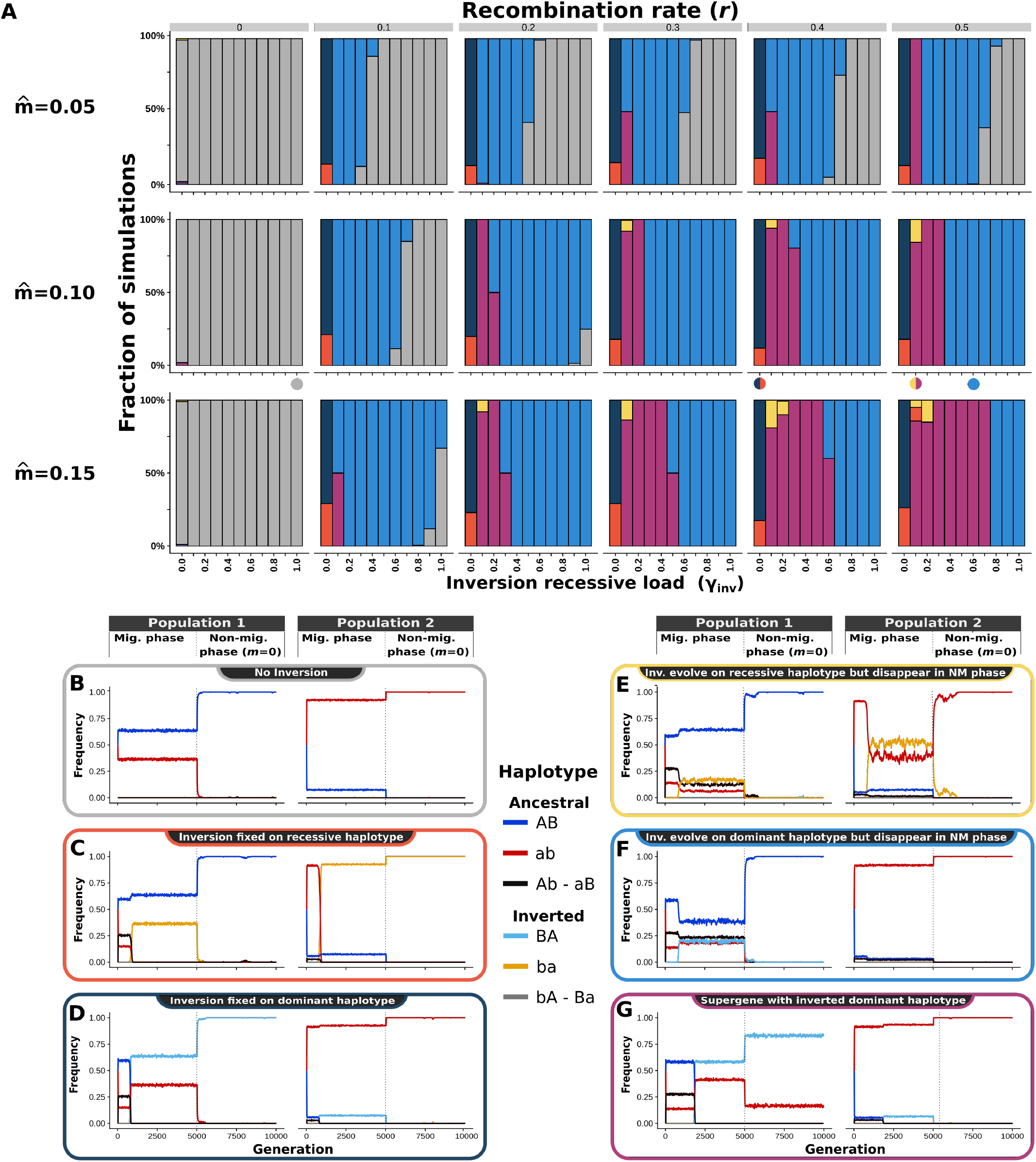
Effect of migration levels on the formation of supergenes. Parameter values used for plot A : *S*_del_=0.8, *S*_div_=0.5, *h*=1.0. Colored dots bellow plot **A** indicate parameters used for plots **B**-**G**.

**Fig. S2.**
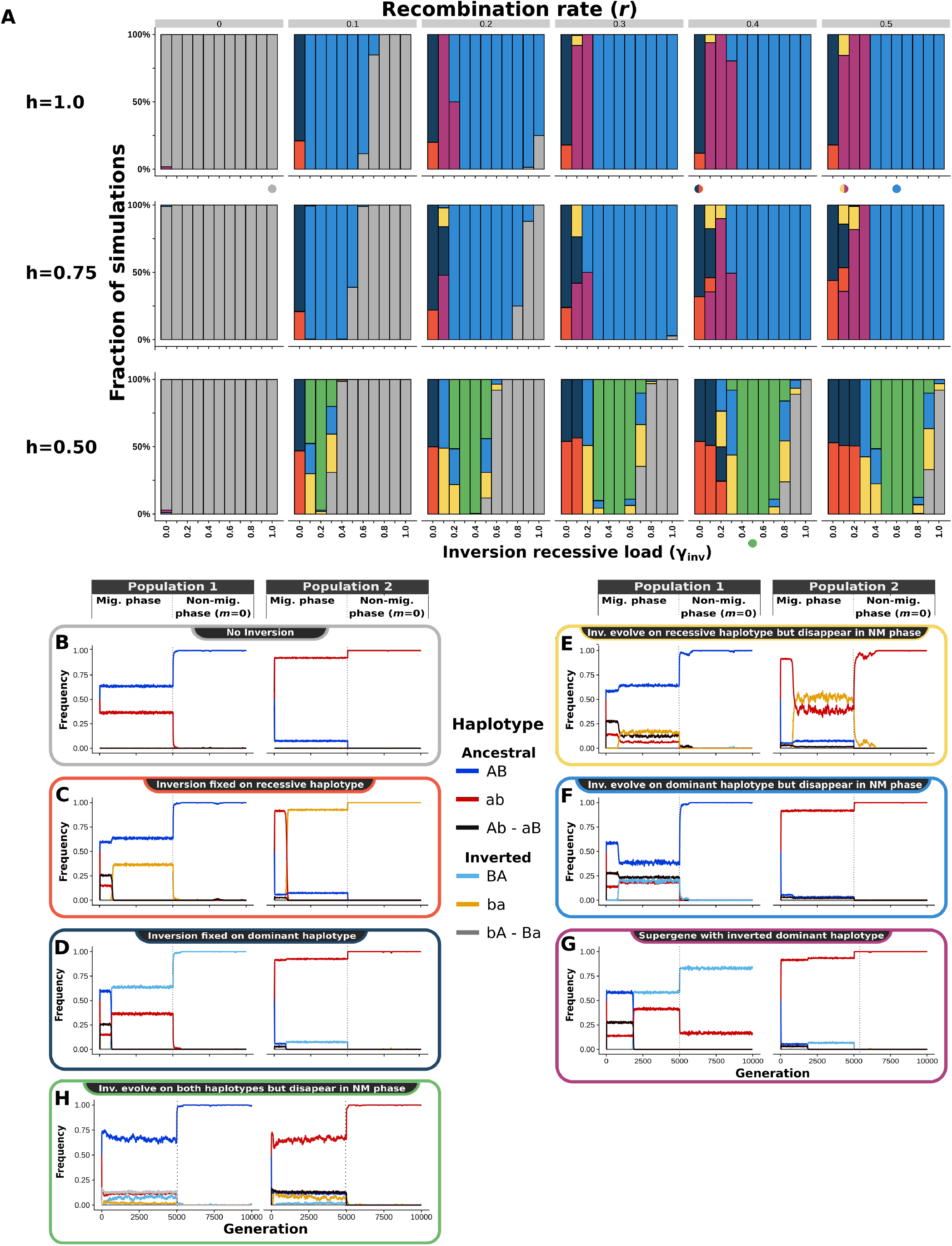
Effect of the dominance levels on the formation of supergenes. Parameter values used for plot A : 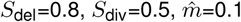 Colored dots bellow plot **A** indicate parameters used for plots **B**-**H**.

**Fig. S3.**
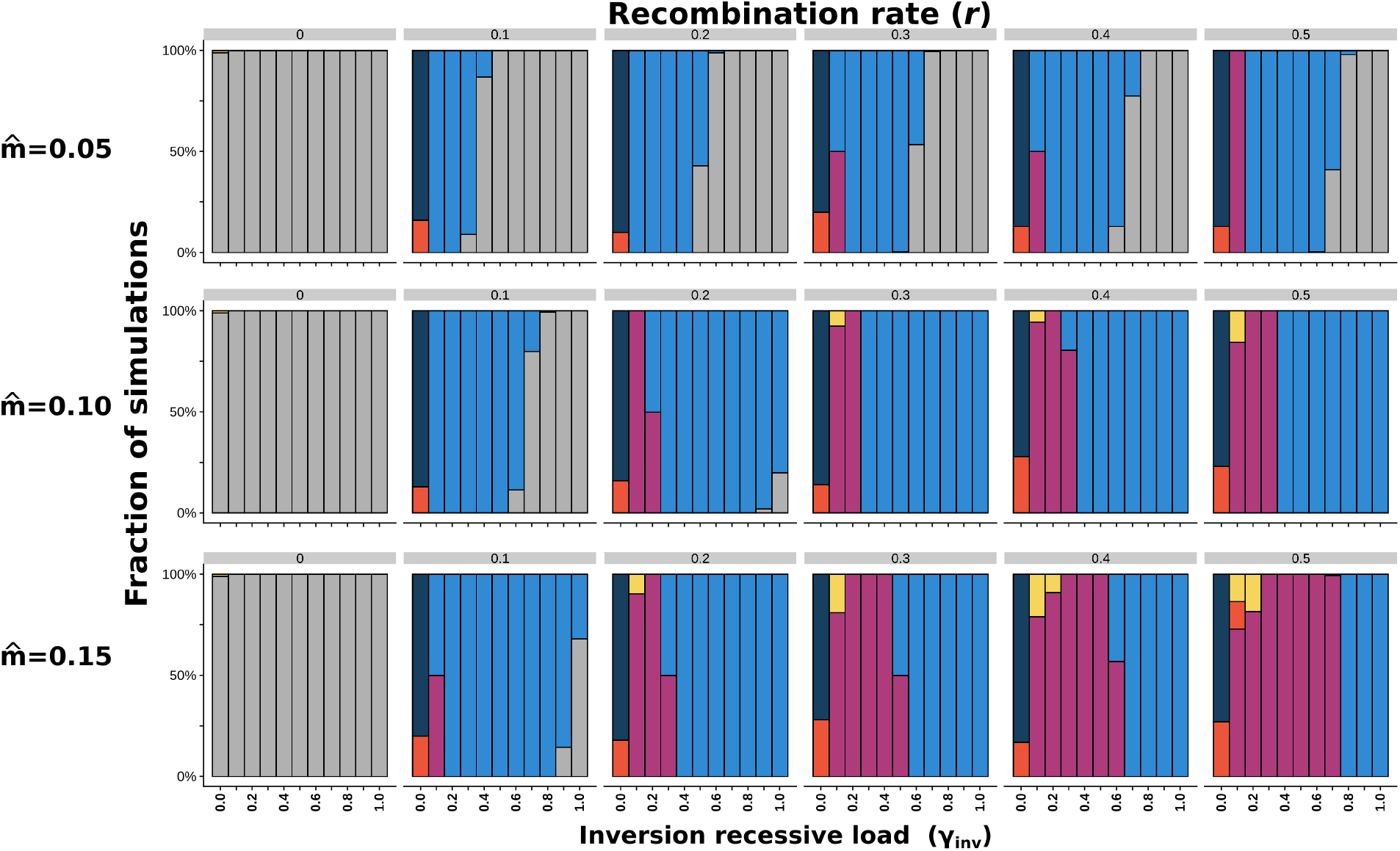
Effect of migration levels on the formation of supergenes with modified life cycle. Life cycle used: Reproduction Selection - Migration. No effect is observed when comparing with the standard life cycle (see Fig. S1). Parameter values used: *S*_del_=0.8, *S*_div_=0.5, *h*=1.0.

**Fig. S4.**
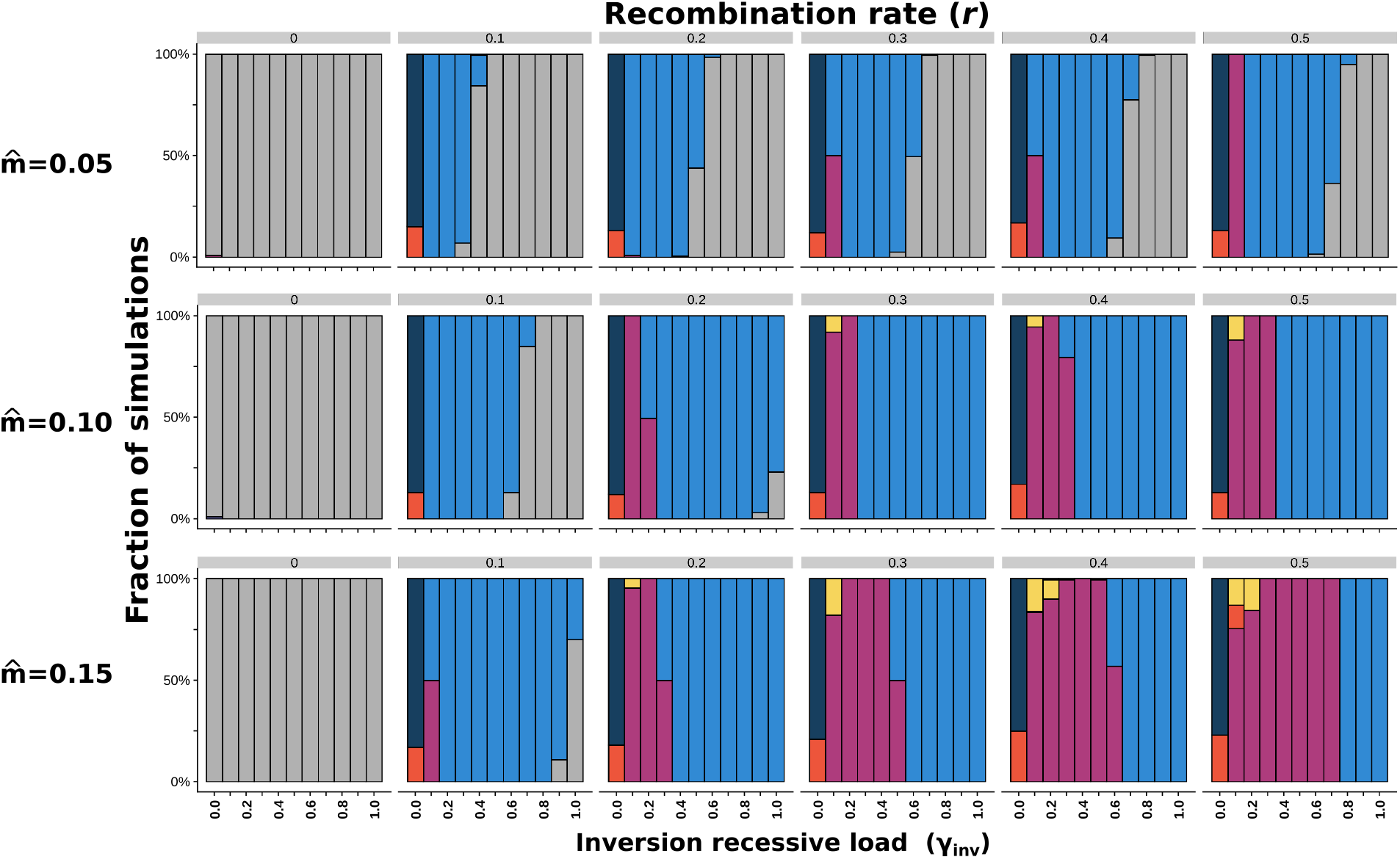
Effect of migration levels on the formation of supergenes with modified life cycle. Life cycle used: Selection - Migration - Reproduction. No effect is observed when comparing with the standard life cycle (see Fig. S1). Parameter values used: *S*_del_=0.8, *S*_div_=0.5, *h*=1.0.

**Fig. S5.**
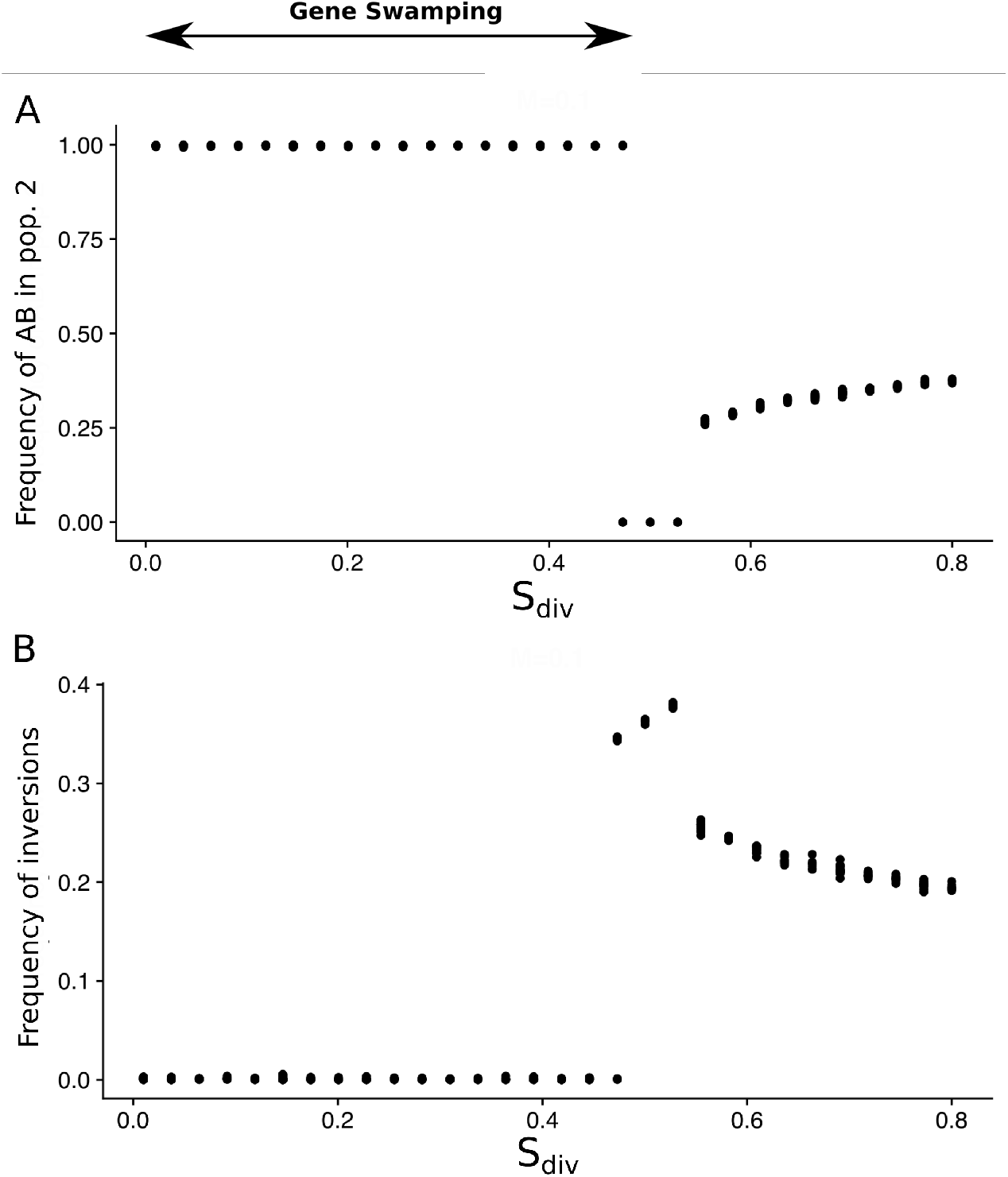
Gene swamping and frequency of inversions in relation to the fitness of immigrant haplotypes. **A** Frequency of haplotype AB in function of *S*_div_. Note : when AB invades population 2, it also invades population 1 (not shown) **B** Frequency of the recombination modifier (inversion) in function of *S*_div_. Parameter values: 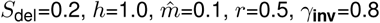.

**Fig. S6.**
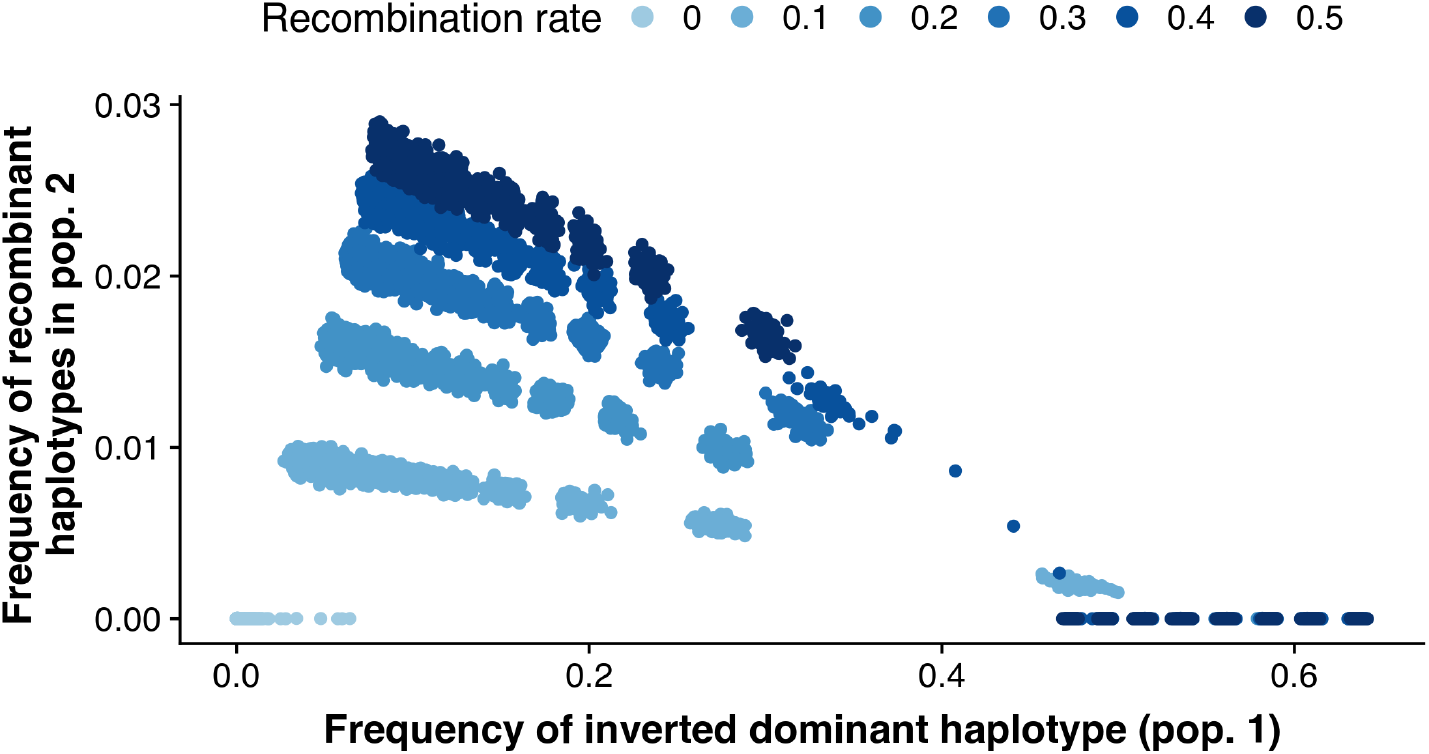
Effect of the spread on an inversion capturing dominant haplotypes (*BA*) on the frequency of recombinants in population 2. Parameter values: *S*_del_=0.2, *S*_div_=0.8, *h*=1.0,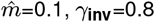.

**Fig. S7.**
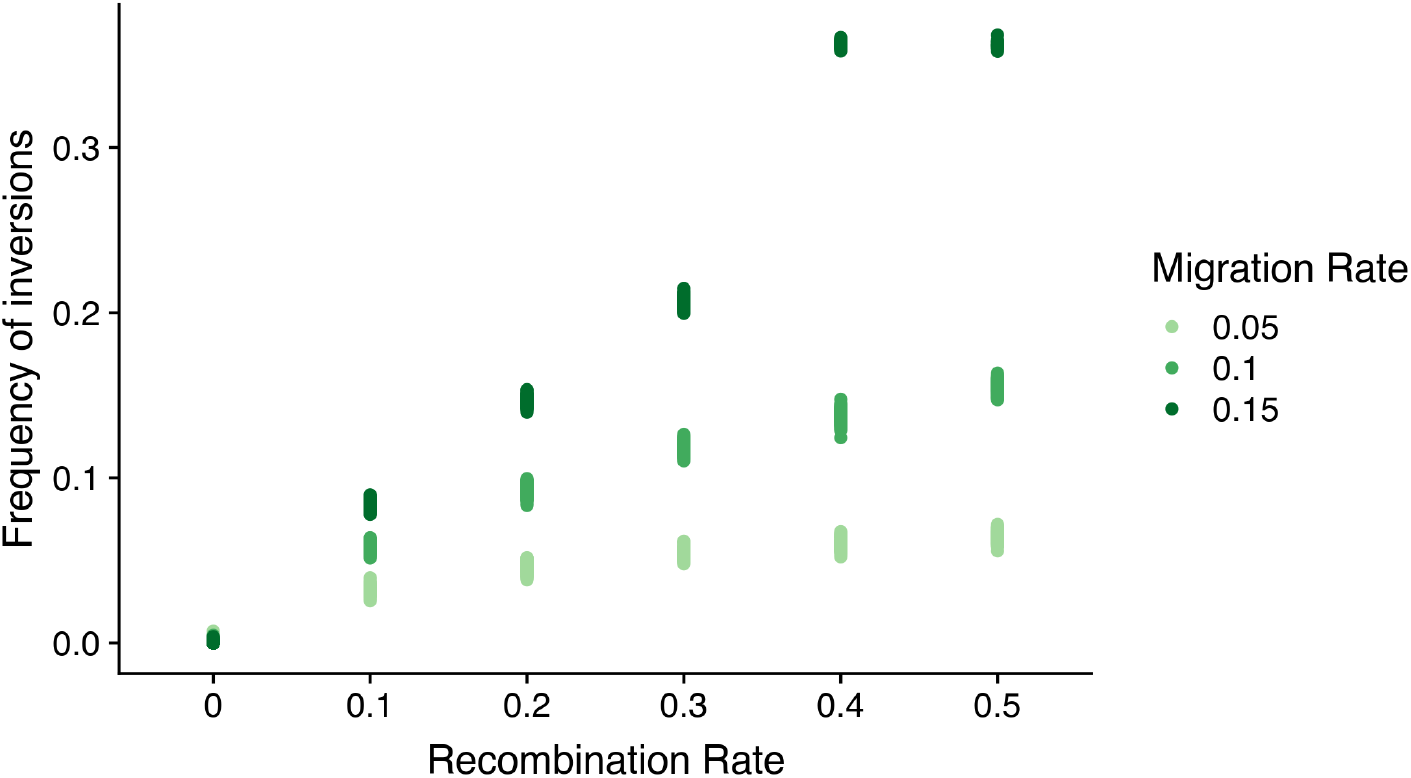
Effect of migration and recombination rate on inversion frequencies during migratory phase. Parameter values: *S*_del_=0.8, *S*_div_=0.5, *r*=0.5, *h*=1,*γ*_**inv**_=0.8.

**Fig. S8.**
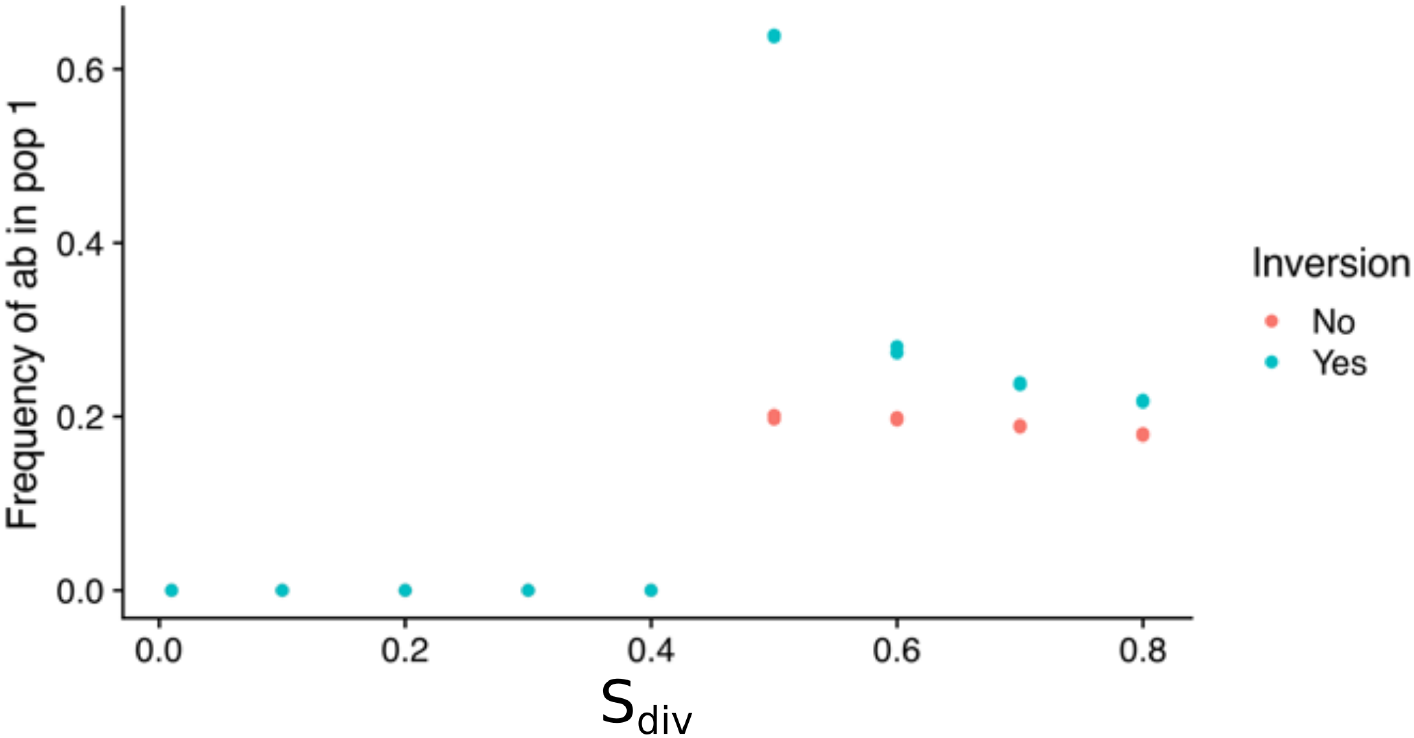
Effects of *S*_div_ (Fitness immigrant) on the frequency of immigrant. Two scenarios are displayed here, depending on whether inversions are allowed to evolve or not. It shows that, as soon as there is no gene swamping (*S*_div_ > 0.4), the evolution of inversions is associated with an increase of immigrant frequency. Parameter values: 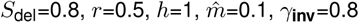.

**Fig. S9.**
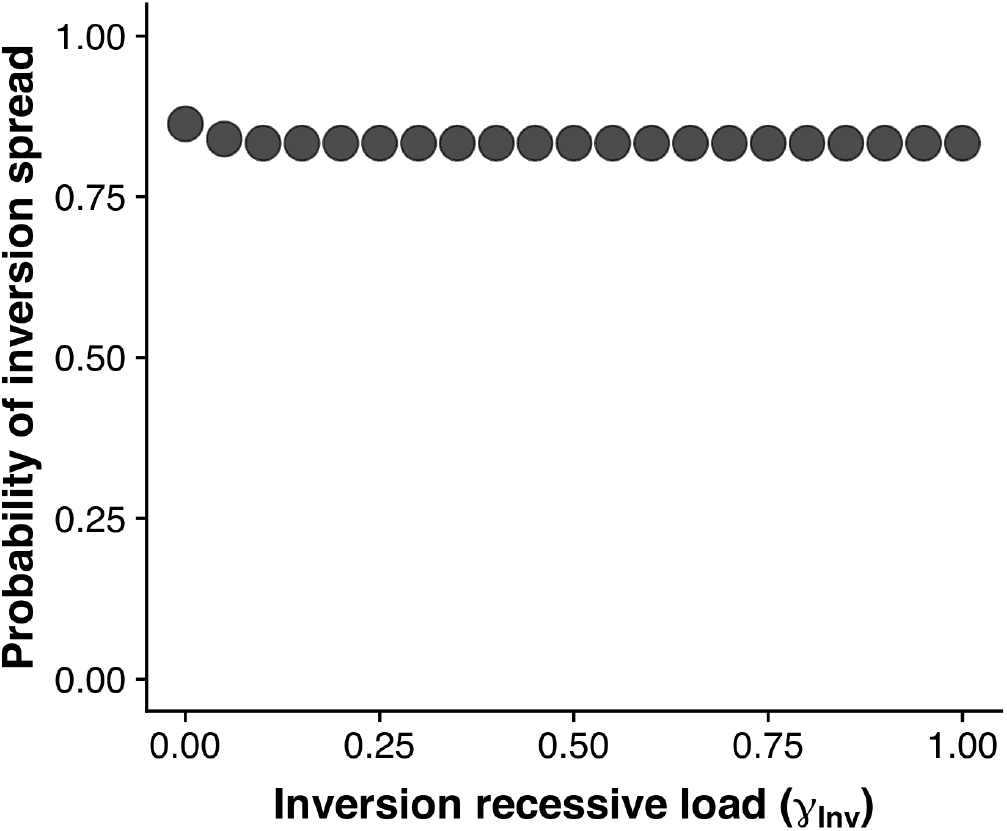
Probability of spread of inversions in function of their recessive load (*γ*_inv_) Probability of inversion frequency > 0.05. Parameter values: *S*_del_=0.8, *S*_div_=0.5, *r*=0.5, m=0.1, *h*=1.

**Fig. S10.**
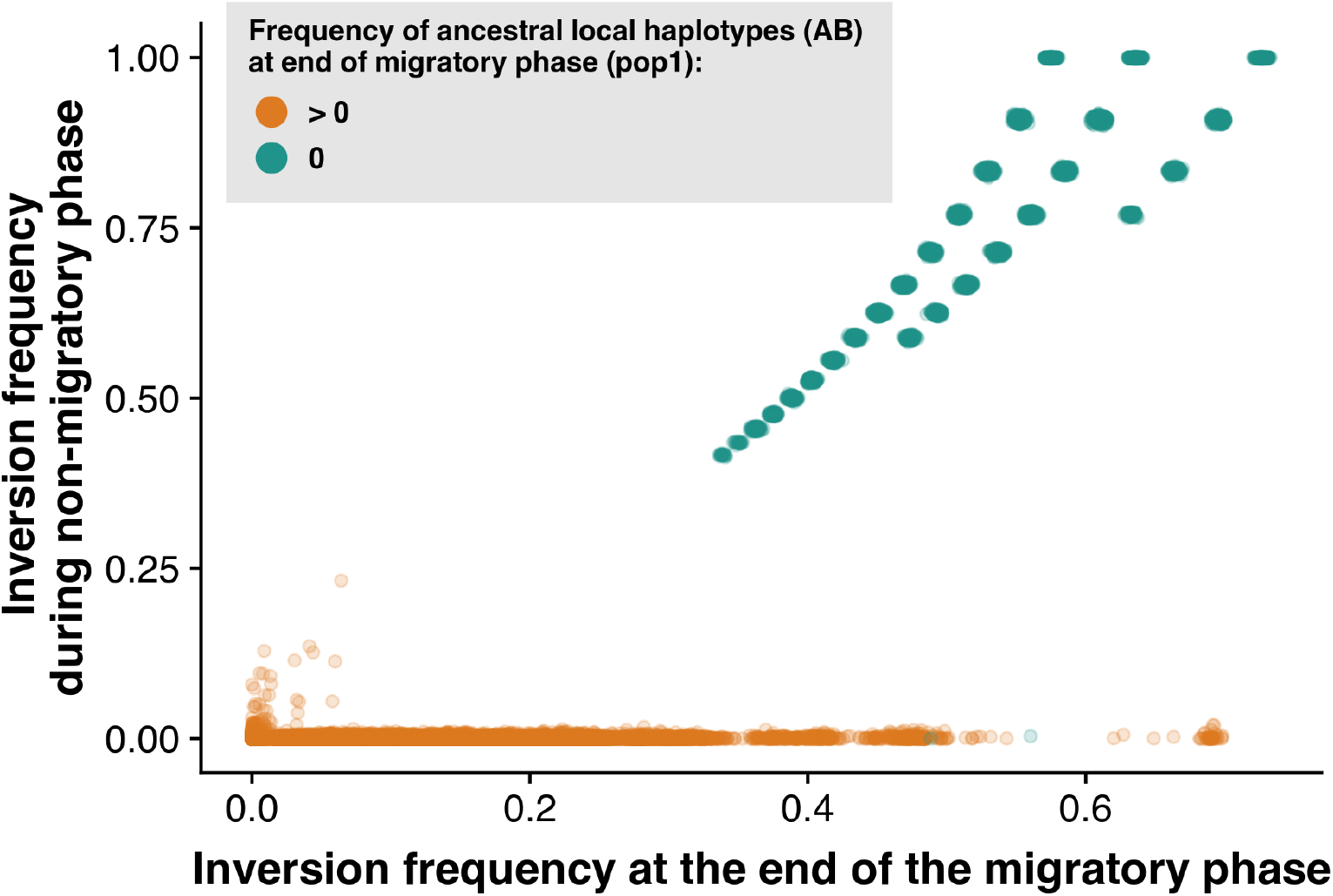
The frequency of the inverted local haplotype in non-migratory phase depends on the frequency of the ancestral local haplotype at the end of the migratory phase. Since inversions are beneficial only when migration occurs between differentiated populations, they can persist during the non-migratory phase only if their ancestral equivalent haplotypes have disappeared during the migratory phase

**Fig. S11.**
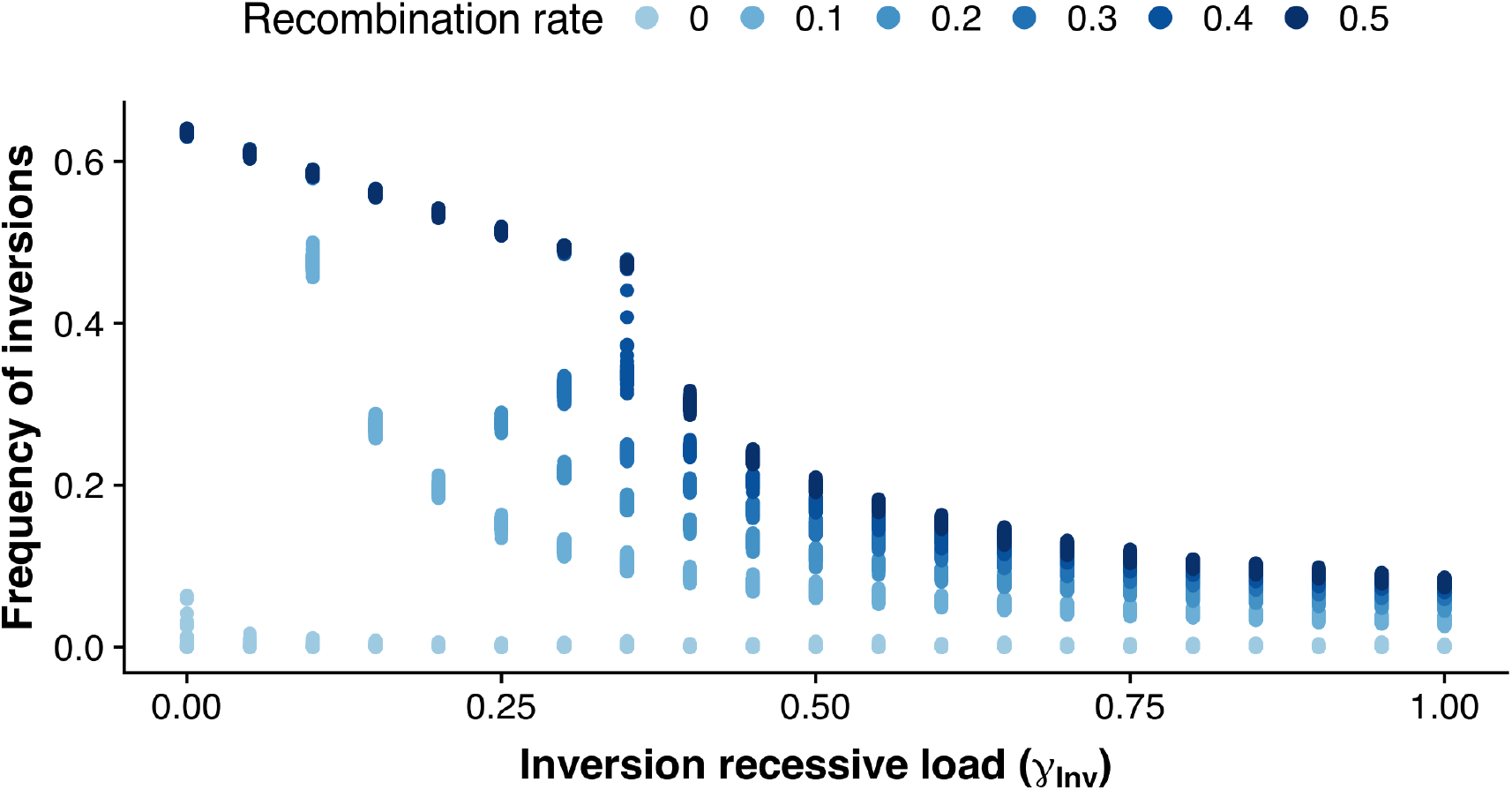
Effect of the inversion load to inversion frequency at end of migratory phase. Parameter values: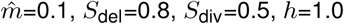.

**Fig. S12.**
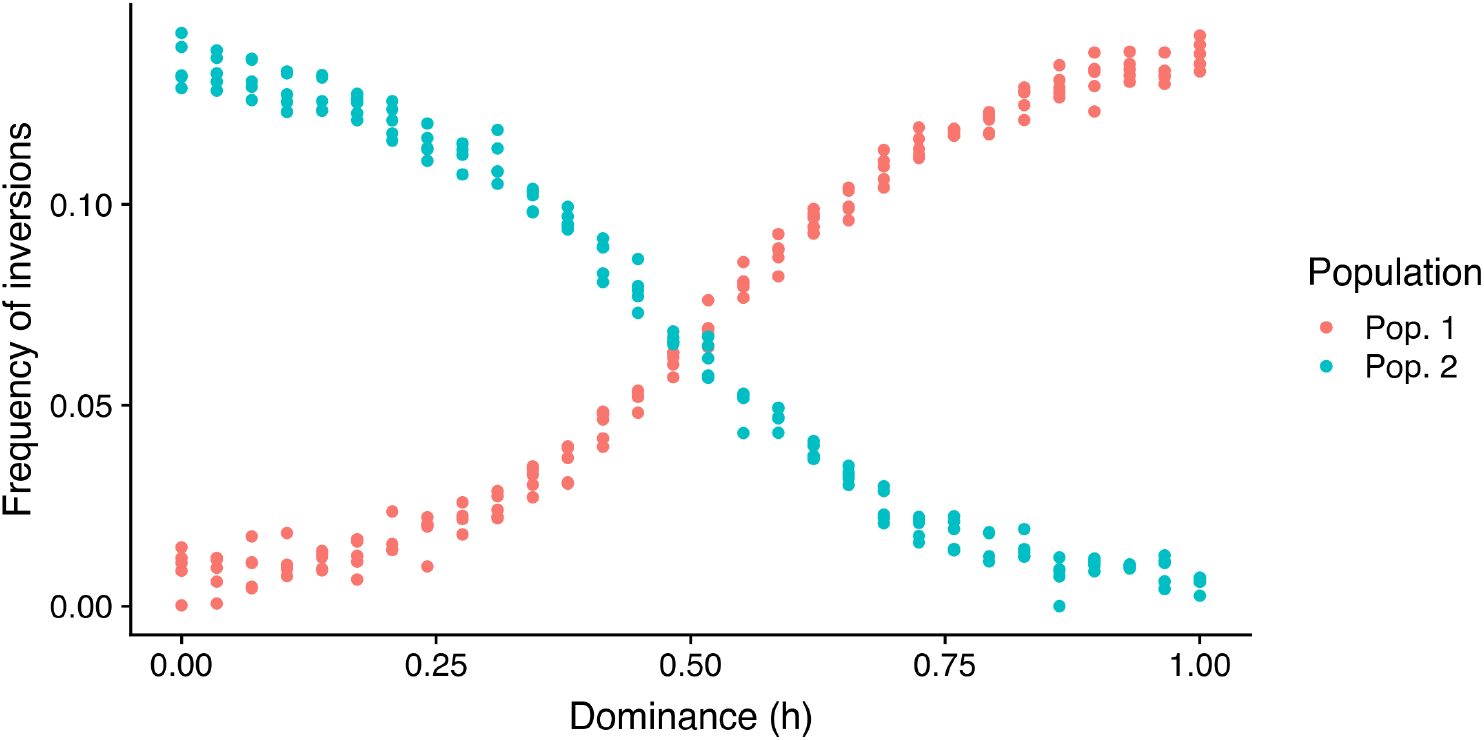
Effect of dominance level on the frequency of inversions in populations. When h<0, it means that the haplotype *ab* is dominant over *AB*. Parameter values: 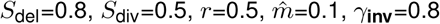.

**Fig. S13.**
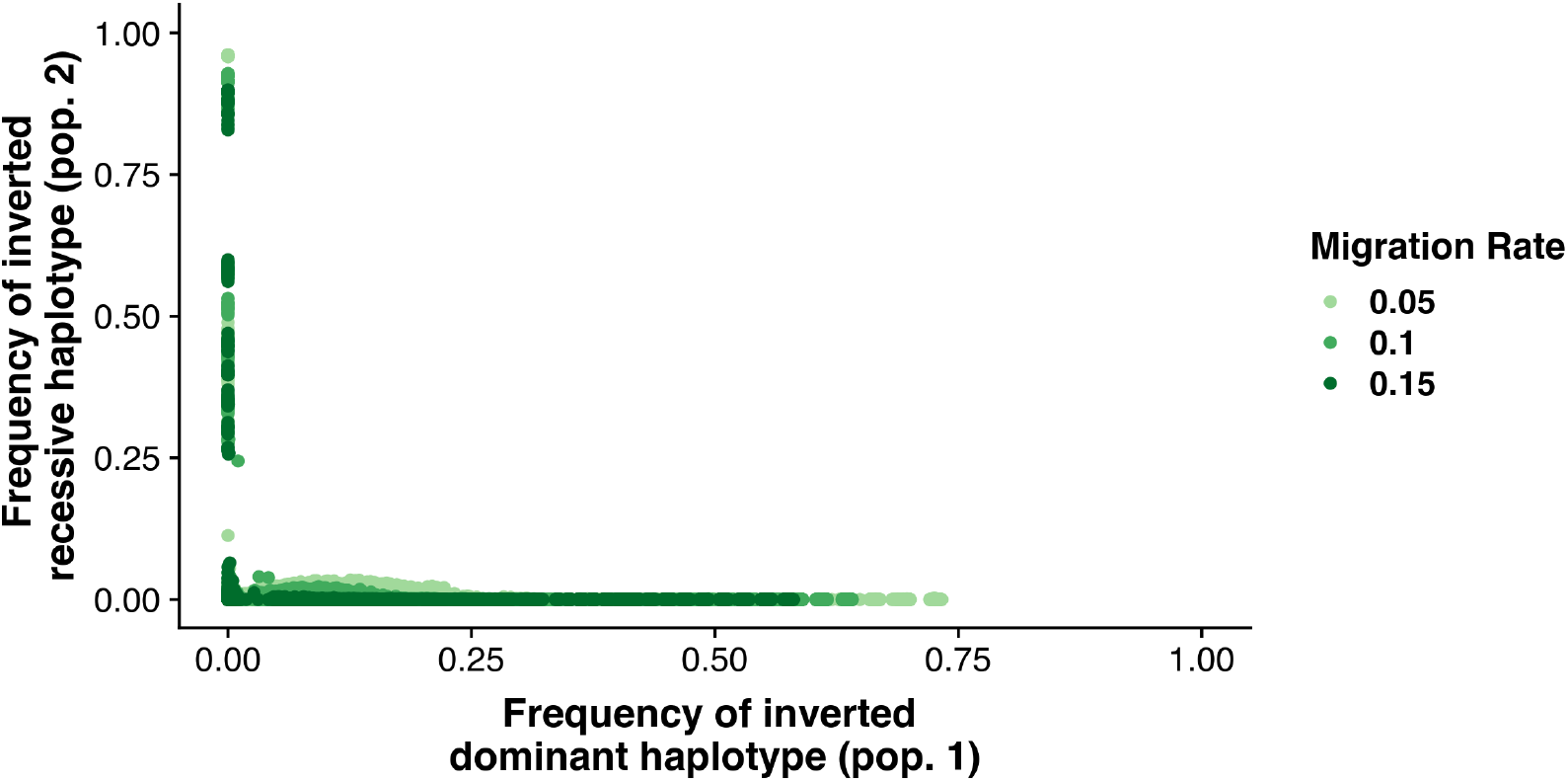
Effect of the spread on an inversion in an haplotype on the spread of inversions in another haplotype. Parameter values: *S*_del_=0.8, *S*_div_=0.5, *r*=0.5, *h*=1, *γ*_**inv**_=8.

**Fig. S14.**
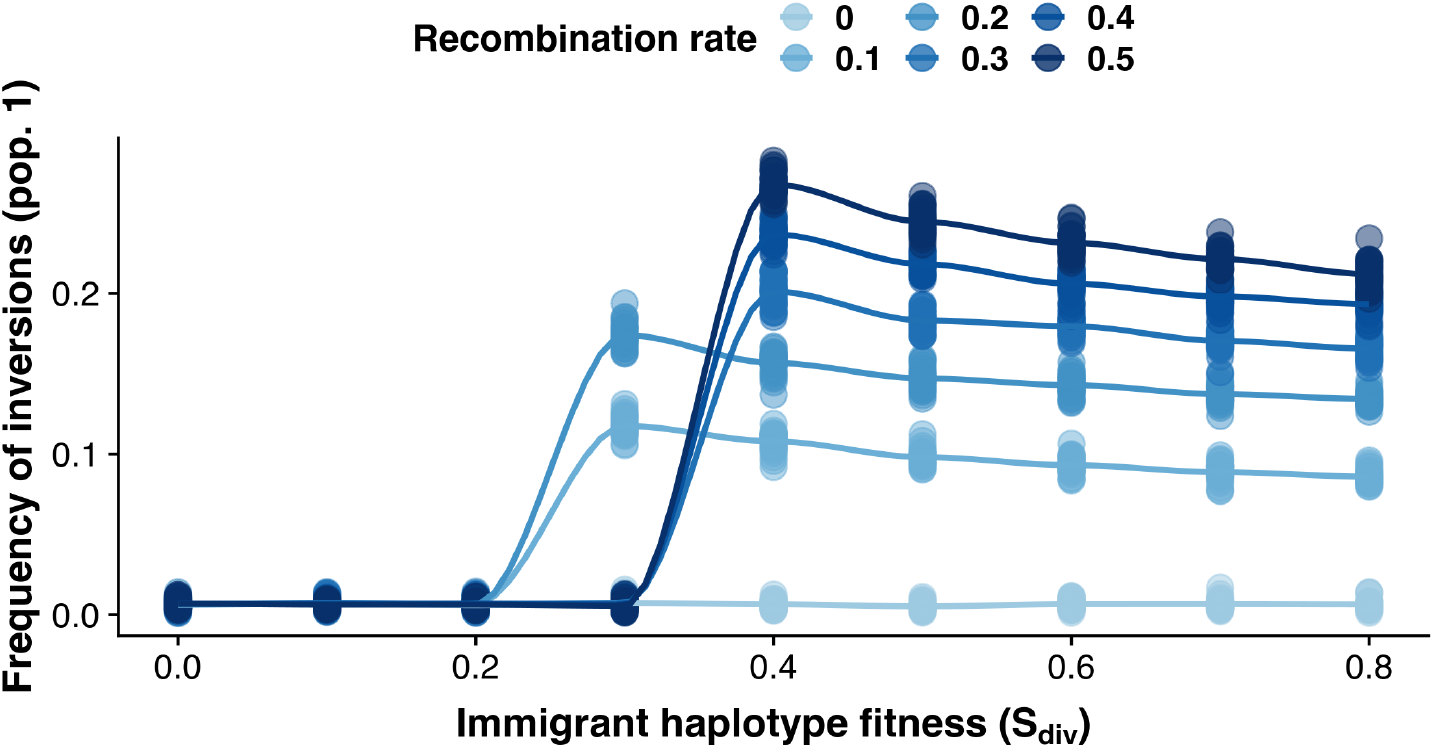
Effects of *S*_div_ (fitness of immigrants) on the frequency of inversions. As soon as there is no gene swamping (*S*_div_ ≤ 0.4), the fitness of immigrants appears as a poor determinant on inversion frequency, contrarily to the recombination rate. Parameter values: *S*_del_=0.8, *r*=0.5, *h*=1,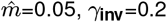.

**Fig. S15.**
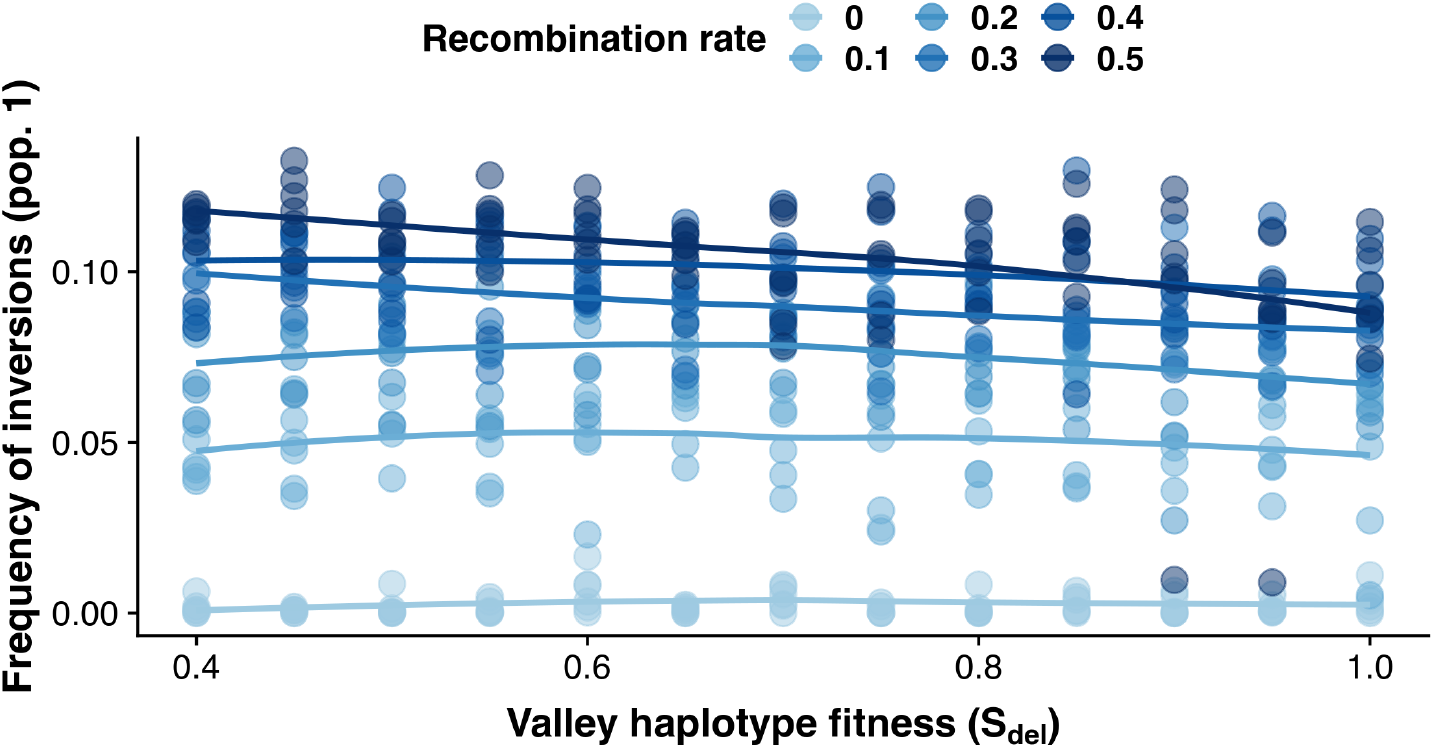
Effects of *S*_del_ (fitness of valley haplotypes) on the frequency of inversions. The fitness of valley haplotypes appears as a poor determinant of inversions frequencies, contrarily to the recombination rate. Parameter values used for these analyses : *S*_div_=0.4, *r*=0.5, *h*=1, 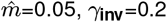.

